# AlphaBeta: Computational inference of epimutation rates and spectra from high-throughput DNA methylation data in plants

**DOI:** 10.1101/862243

**Authors:** Yadollah Shahryary, Aikaterini Symeonidi, Rashmi R. Hazarika, Johanna Denkena, Talha Mubeen, Brigitte Hofmeister, Thomas van Gurp, Maria Colomé-Tatché, Koen Verhoeven, Gerald Tuskan, Robert J Schmitz, Frank Johannes

## Abstract

**Introduction:** Heritable changes in cytosine methylation can arise stochastically in plant genomes independently of DNA sequence alterations. These so-called ‘spontaneous epimutations’ appear to be a byproduct of imperfect DNA methylation maintenance during mitotic or meitotic cell divisions. Accurate estimates of the rate and spectrum of these stochastic events are necessary to be able to quantify how epimutational processes shape methylome diversity in the context of plant evolution, development and aging.

**Method:** Here we describe *AlphaBeta*, a computational method for estimating epimutation rates and spectra from pedigree-based high-throughput DNA methylation data. The approach requires that the topology of the pedigree is known, which is typically the case in the experimental construction of mutation accumulation lines (MA-lines) in sexually or clonally reproducing species. However, this method also works for inferring somatic epimutation rates in long-lived perennials, such as trees, using leaf methylomes and coring data as input. In this case, we treat the tree branching structure as an intra-organismal phylogeny of somatic lineages and leverage information about the epimutational history of each branch.

**Results:** To illustrate the method, we applied *AlphaBeta* to multi-generational data from selfing- and asexually-derived MA-lines in Arabidopsis and dandelion, as well as to intra-generational leaf methylome data of a single poplar tree. Our results show that the epimutation landscape in plants is deeply conserved across angiosperm species, and that heritable epimutations originate mainly during somatic development, rather than from DNA methylation reinforcement errors during sexual reproduction. Finally, we also provide the first evidence that DNA methylation data, in conjunction with statistical epimutation models, can be used as a molecular clock for age-dating trees.

**Conclusion:** *AlphaBeta* faciliates unprecedented quantitative insights into epimutational processes in a wide range of plant systems. Software implementing our method is available as a Bioconductor R package at http://bioconductor.org/packages/3.10/bioc/html/AlphaBeta.html

## Introduction

Cytosine methylation is an important chromatin modification and a pervasive feature of most plant genomes. It has major roles in the silencing of transposable elements (TEs) and repeat sequences, and is also involved in the regulation of some genes [1]. Plants methylate cytosines at symmetrical CG and CHG sites, but also extensively at asymmetrical CHH sites, where H= A, T, C. The molecular pathways that establish and maintain methylation in these three sequence contexts are well-characterized [2], and are broadly conserved across plant species [3] [4] [5] [6] [7]. Despite its tight regulation, the methylation status of individual cytosines or of clusters of cytosines is not always faithfully maintained across cell divisions. As a result, cytosine methylation is sometimes gained or lost in a stochastic fashion, a phenomenon that has been termed ‘spontaneous epimutation’. In both animals and plants, spontaneous epimutations have been shown to accumulate throughout development and aging [8] (see co-submission), probably as a byproduct of the mitotic replication of small stem cell pools that generate and maintain somatic tissues (**Fig. 1A**).

**Figure 1:**
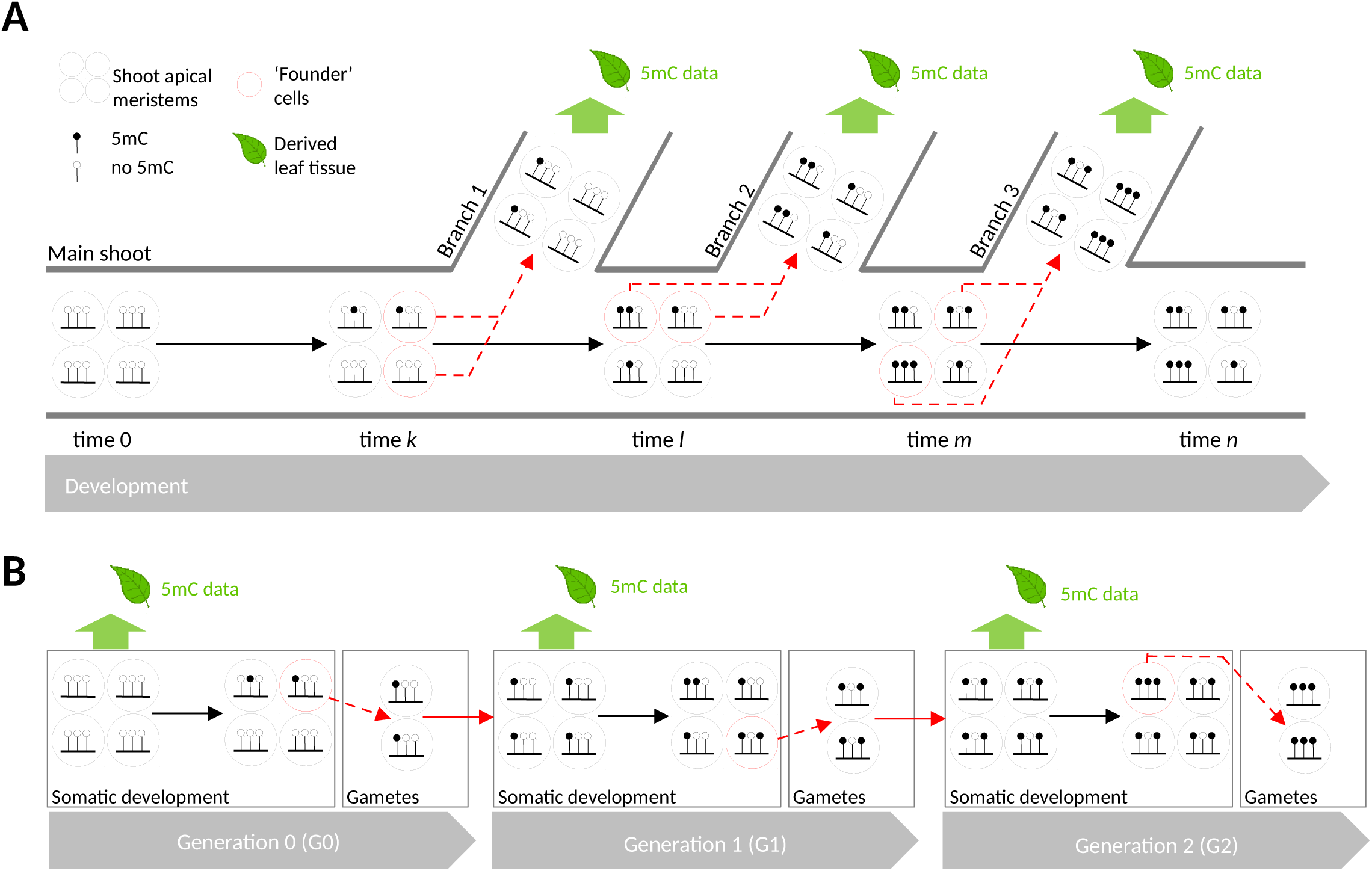
Schematic of the developmental origin of somatic and ‘germline’ epimutations in plants. **A.** The failure to maintain the methylation status of cytosines during the mitotic maintenance of shoot apical meristematic cell pools leads to spontaneous somatic epimutations. Shown here are only spontaneous gains of methylation, for simplicity. A small set of ‘founder’ cells gives rise to lateral branches at developmental times *k, l*, and *m*. The random sampling of these founder cells can create a bottleneck which increases the frequency of somatic epimutations in the cell populations of lateral branches. Somatic epimutation accumulation in shoot apical meristems thus leads to increased 5mC divergence between leafs originating from different lateral branches (e.g. leaf methylomes from Branch 1 and 2 are more similar than those from Branch 1 and 3). **B.** Shown is the origin of ‘germline’ epimutations. We assume that the male and female sexual cell lineages are derived from somatic precursors late in development. Somatically-aquired epimutations can thus pass to the gametes. Combined with a lack of proper reprogramming, these somatic epimutations can be inherited to subsequent generations. The accumulation of these ‘germline’ epimutations leads to increased methylome divergence between leaves sampled from different generations (e.g. leaf methylomes from generation 0 and 1 are more similar than those from generation 0 and 2).

However, in plants, spontaneous epimutations are not only confined to somatic cells, but occasionally pass through the gametes to subsequent generations [9] [10](**Fig. 1B**). In the model plant *Arabidopsis thaliana* (*A. thaliana*) these transgenerationally heritable (i.e ‘germline’) epimutations are mainly restricted to CG sites, and appear to be absent or not detectable at CHG and CHH sites [11]. Estimates in *A. thaliana* indicate CG ‘germline’ epimutations are about five orders of magnitude more frequent than genetic mutations (∼ 10^−4^ vs. ∼ 10^−9^ per site per haploid genome per generation) [12] [13]. Because of these relatively high rates, CG methylation differences accumulate rapidly in the *A. thaliana* genome, and generate substantial methylation diversity among individuals in the course of only a few generations [13] [14] [15].

A key experimental challenge in studying epimutational processes in a multi-generational setting is to be able to distinguish ‘germline’ epimutations from other types of methylation changes, such as those that are associated with segregating genetic variation or transient environmental perturbations [16]. Mutation Accumulation (MA) lines grown in controlled laboratory conditions are a powerful experimental system to achieve this. MA lines are derived from a single isogenic founder and are independently propagated for a large number of generations. The lines can be advanced either clonally or sexually, i.e. self-fertilization or sibling mating (**Fig. 2A**). In clonally produced MA lines the isogenicity of the founder is not required because the genome is ‘fixed’ due to the lack of genetic segregation.

**Figure 2:**
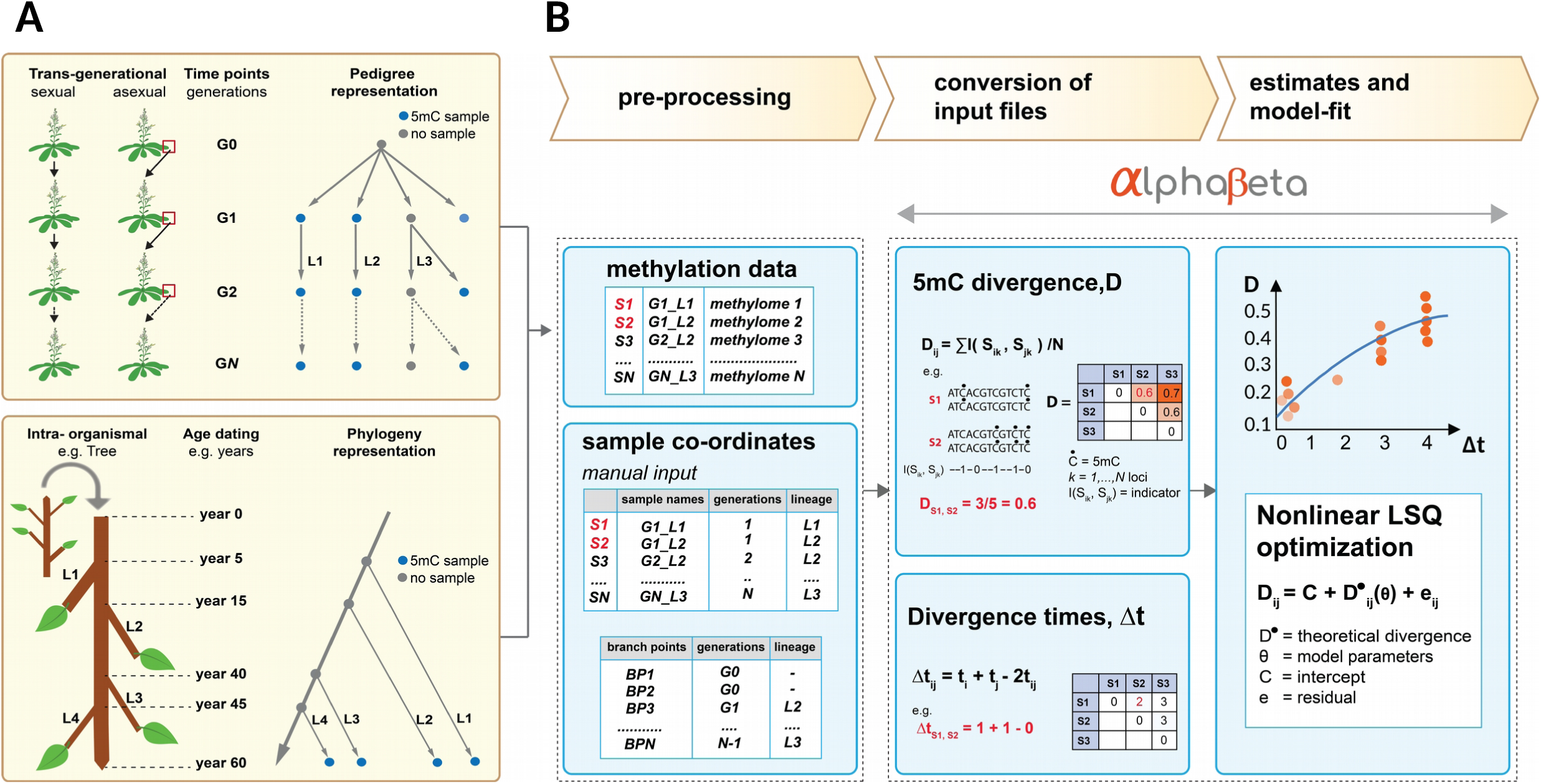
Overview of the *AlphaBeta* computational pipeline. **A.** *Top panel:* Experimental construction of multi-generational (G0 to G*N*) Mutation Accumulation (MA) lines through sexual (selfing or sibling mating) or asexual (clonal) propagation. The relationship between the different lineages (L1 to L3) can be represented as a pedigree. The pedigree branch point times and the branch lengths are typically known, *a priori*, from the experimental design. 5mC sampling can be performed at selected generations, either from plant material of direct progenitors or from siblings of those progenitors. The data can be used to estimate the rate and spectrum of ‘germline’ epimutations. Bottom panel: Long-lived perennials, such as trees, can be viewed as a natural mutation accumulation system. In this case, the tree branching structure can be treated as an intra-organismal phylogeny of somatic lineages that carry information about the epimutational history of each branch. 5cC samples can be performed on leaf tissues from selected branches. Along with coring data, the leaf methylomes can be used to estimate the rate and spectrum of somatic epimutations. **B.** Data pre-processing: *AlphaBeta* requires three input files: 1. cytosine methylation state calls for each sample; 2. coordinates of each sample within the pedigree; 3. coordinates of the branch points (or nodes) of the pedigree/phylogeny. File conversion: Using the input files, *AlphaBeta* calculates the 5mC divergence (D) as well as divergence time (Δt) between all sample pairs. Model estimation: *AlphaBeta* features a number of competing epimutation models that model the relationship between D and Δt. The model parameters are estimated using numerical non-linear least squares optimization. Competing models can be formally compared, and allow for tests of selection and neutrality.

The kinship among the different MA lineages can be presented as a pedigree (**Fig. 2A**). The structure (or topology) of these pedigrees is typically known, *a priori*, as the branch-point times and the branch lengths are deliberately chosen as part of the experimental design. In conjunction with multi-generational methylome measurements MA lines therefore permit ‘real-time’ observations of ‘germline’ epimutations against a nearly invariant genomic background, and can facilitate estimates of the per generation epimutation rates [11]. Sequenced methylomes from a large number of sexually-derived MA lines are currently available in *A. thaliana* [17] [12] [18] [13] [19] [15], rice [20], and various other MA-lines are currently under construction for epimutation analysis in different genotypes, enviornmental conditions and plant species.

Beyond experimentally-derived MA lines, natural mutation accumulation systems can also be found in the context of plant development and aging. An instructive example are long-lived perennials, such as trees, whose branching structure can be interpreted as a pedigree (or phylogeny) of somatic lineages that carry information about the epimutational history of each branch [21]. In this case, the branch-point times and the branch lengths can be determined *ad hoc* using coring data, or other types of dating methods (**Fig. 2A**). By combining this information with contemporary leaf methylome measurements it is possible to infer the rate of somatic epimutations as a function of age (see co-submission).

Attempts to infer the rate of spontaneous epimutations in these diverse plant systems are severely hampered by the lack of available analytical tools. Naive approaches that try to count the number of epimutations per some unit of time cannot be used in this setting, because DNA methylation measurements are far too noisy. On the technological side, this noise stems from increased sequencing and alignment errors of bisulphite reads and bisulphite conversion inefficiencies. On the biological side, increased measurement error may result from within-tissue heterogeneity in 5mC patterns [22] and the fact that DNA methylomes are in part transcriptionally responsive to variation in enviornmental/laboratory conditions [23]. To overcome these challenges, we previously implemented a model-based estimation method, which was originally designed for the analysis of selfing-derived mutation accumulation lines [13]. This approach appropriately accounts for measurement error in the data by describing the time-dependent accumulation of epimutations through an explicit statistical model (**Fig. 2B**). Fitting this model to pedigree-based 5mC measurements yields estimates of the rate of spontaneous methylation gains and losses, and provides a quantitative basis for predicting DNA methylation dynamics over time.

Here, we build on this method and present *AlphaBeta*, the first comprehensive software package for infering the rate and spectrum of ‘germline’ and somatic epimutations in plants. *AlphaBeta* can be widely applied to multi-generational data from sexually- or clonally-derived MA lines, as well as to intra-generational data from long-lived perennials such as trees. Drawing on novel and published data, we demonstrate the power and versatility of our approach, and make recommendations regarding its implementation.

## Results

### Conceptual overview of the method

We start from the assumption that 5mC measurements have been obtained from multiple sampling time-points throughout the predigree (**Fig. 2A**). These measurements can come from whole genome bisulphite sequencing (WGBS) [24] [25], reduced representation bisulphite sequencing (RRBS) [26], or epigenotyping by sequencing (epiGBS) [27] technologies, and possibly also from array-based methods. We only require that a ‘sufficiently large’ number of loci has been measured. Moreover, with multigenerational data we allow measurements to come from plant material of direct progenitors, or else from individual or pooled siblings of those progenitors (**Fig. 2A**).

For the *i*th sequenced sample, we let *s*_*ik*_ be the observed methylation state at the *k*th locus (*k* = 1 … *N*). Here, the *N* loci can be individual cytosines or pre-defined regions (i.e. cluster of cytosines). We assume that *s*_*ik*_ takes values 1, 0.5 or 0, according to whether the diploid epigenotype at that locus is *m/m, m/u, u/u*, respectively, where *m* is a methlylated and *u* is an unmethylated epiallele. Using this coding, we calculate the total 5mC divergence, *D*, between any two samples *i* and *j* as follows:

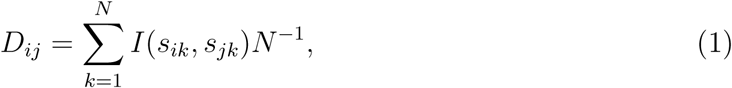

where *I*(·) is an indicator function, such that *I*(*s*_*ik*_, *s*_*jk*_) is equal to 0 if *s*_*ik*_ = *s*_*jk*_, 0.5 if *s*_*ik*_ = 0.5 and *s*_*jk*_ ∈ {0, 1}, 0.5 if *s*_*jk*_ = 0.5 and *s*_*ik*_ ∈ {0, 1}, and 1 if *s*_*jk*_ = 1 and *s*_*ik*_ = 0. We suppose that *D*_*ij*_ is related to the divergence time (Δt) of samples *i* and *j* through an underlying epimutation model *M*_Θ_ (**Fig. 2B**). The software automatically calculates *D*_*ij*_ and Δt for all unique sample pairs using as input the methylation state calls and the pedigree coordinates of each sample (**Fig. 2B**).

We model the 5mC divergence using

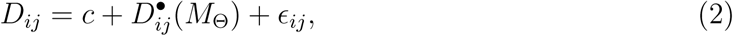

where *ϵ*_*ij*_ ∼ *N* (0, *σ*^2^) is the normally distributed residual error, *c* is the intercept, and 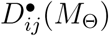 is the expected divergence as a function of epimutation model *M* with parameter vector Θ. Parameter vector Θ contains the unknown spontaneous gain rate *α*, the loss rate *β*, selection coefficient *w*, as well as the unknown proportion *γ* of epiheterozygote loci in the most recent common founder of samples *i* and *j*. Model *M* can take four different forms, which we denote by ABneutral, ABmm, ABuu, ABnull. Model ABneutral assumes that spontaneous 5mC gains and losses accumulate neutrally over time, ABmm assumes that the accumulation is partly shaped by selection against spontaneous losses of 5mC, ABuu assumes that the accumulation is partly shaped by selection against spontaneous gains, and ABnull is the null model of no accumulation (see Methods). Each model is specified in such a way as to reflect the particular mating system that was used to generate the pedigree (e.g. selfing, clonal, somatic), and can be applied to any arbitrary pedigree structure (i.e. topology) as long as the lineage branch points and the branch lengths are known. As mentioned above, this latter information is typically known *a priori* in the context of experimentally-derived MA lines or it can be supplied *ad hoc*, for example from coring data as in the case of trees.

To obtain estimates for Θ we seek to minimize

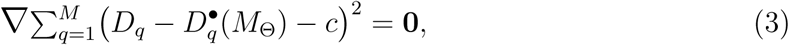

where the summation is over all *M* unique pairs of sequenced samples in the pedigree. The minimization of this equation is a problem in non-linear least square regression. The theoretical derivation of 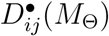 and details regarding parameter estimation are provided in the Methods section.

### Analysis of spontaneous epimutations in selfing-derived *A. thaliana* MA-lines

To illustrate our method, we first analyzed three *A. thaliana* MA pedigrees (MA1_1, MA1_3, MA3, see **Fig. 3A**). We chose these MA pedigrees because they differ markedly in their topologies, 5mC sampling strategies, sequencing method and depth (**Fig. 3A-B, Table S1**). All MA pedigrees were derived from a single Col-0 founder accession. The first MA pedigree (MA1_1) was origially published by Becker et al. [17]. The pedigree data consists of 11 independent lineages with sparsely collected WGBS samples (∼ 19.2X coverage) from generations 3, 31 and 32, and a maximum divergence time (Δt) of 64 generations. MA1_3 was previously published by van der Graaf et al. [13]. This data consists of single lineage with dense MethylC-seq measurements (∼ 13.8X coverage) from generations 18 to 30, and a maximum Δt of 13 generations. Finally, we present a new pedigree (MA3), which consists of 2 lineages with dense MethylC-seq measurements (∼ 20.8X coverage) from generations 0 to 11, and a maxium Δt of 22 generations. Unlike MA1_1 and MA1_3, MA3 has 5mC measurements from progenitor plants of each sampled generation, rather than from siblings of those progenitors (**Fig. 3A**). Further information regarding the samples, sequencing depths and platforms is provided in **Table S1**. A detailed description of data pre-processing and methylation state calling can be found in the Methods section.

**Figure 3:**
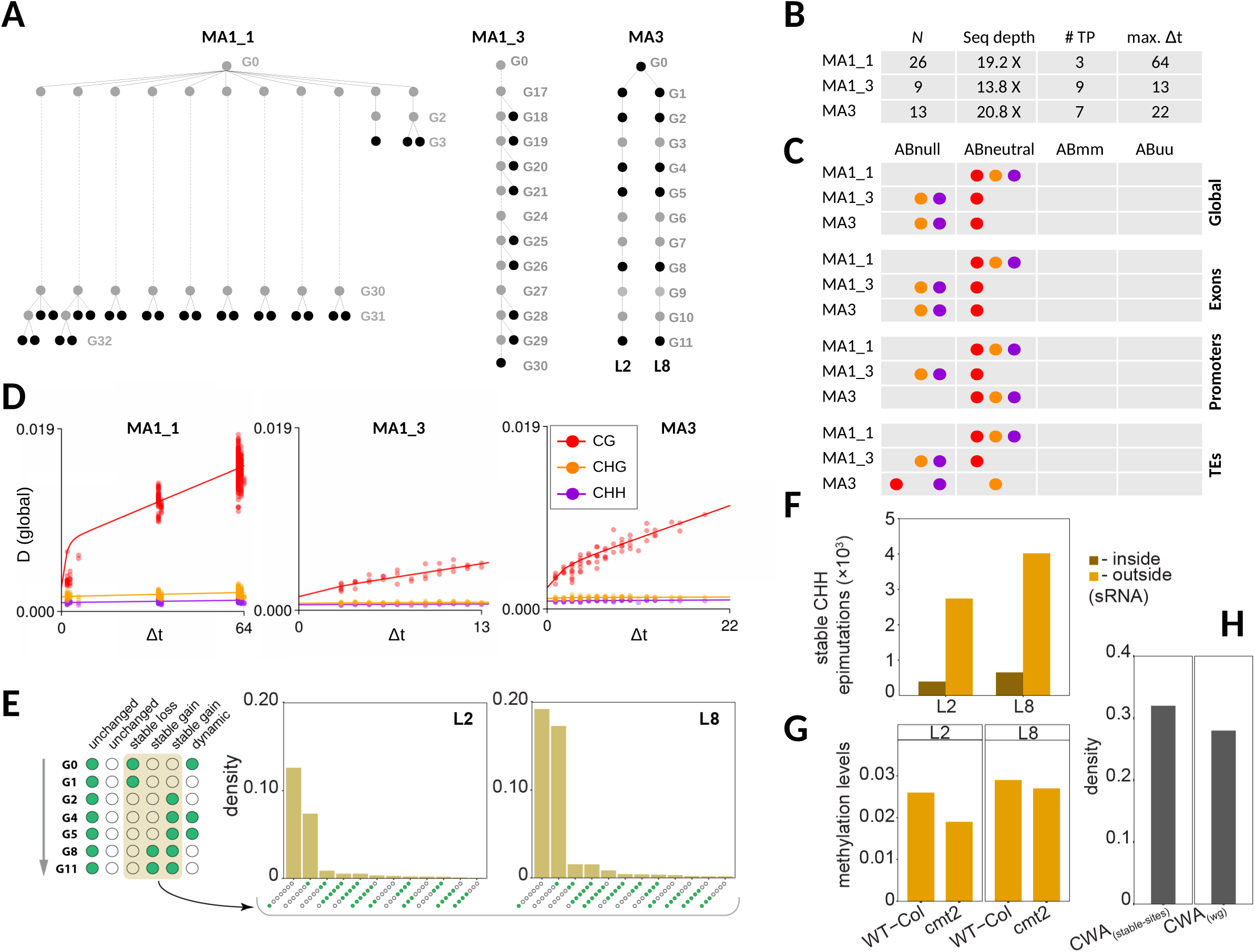
Analysis of ‘germline’ epimutations in *A. thaliana* Mutation Accumulation (MA) lines. **A.** Three different MA-pedigrees were analyzed. All three pedigree were derived from a single Columbia (Col-0) inbred genotype. Two of the pedigrees were previously published (MA1_1, Becker *et al.* 2011; MA1_3, van der Graaf *et al*. 2015), and one pedigree (MA3) is new. These three MA pedigrees were chosen because they differ in their typologies, 5mC measurement strategies, and the temporal resolution of the 5mC samples. **B.** Overview of the data: *N* is the total number of sequenced samples; Seq depth is the average sequence depths of the samples; # TP is the number of unique time-points (or generations) that are sampled; max. Δt is the maximum divergence time (in generations) in the pedigree. **C.** We applied four competing models to these pedigree data (Abnull, Abneutral, ABmm and Abuu). These models are described in the main text. The best fitting model is indicated for each MA-pedigree, sequence context (CG, CHG and CHH) and genomic feature (global, exons, promoters, TEs). We detected consistent and significant accumulation of 5mC divergence (*D*) over generation time (Δt) for context CG, both at the global and within specific genomic features. Model inference for CHG and CHH was more variable and dependent the size of the dataset, both in terms of the number of samples as well as on the maximum divergence time. **D.** Shown are the fits of the best fitting models for each pedigree and context. **E.** Schematic representation of transgenerational CHH epimutations. The barplots indicate the density of stable CHH epimutations in lineages L2 and L8 of the MA3 pedigree. **F.** CHH sites featuring stable epimutations tend to fall outside of sRNA clusters in lineages L2 and L8. **G.** Analysis of *cmt2* mutant and Col-0 wt from Stroud et al. 2013 show loss of methylation in the mutant at the stable CHH epimutation sites, indicating that these loci are targeted by CMT2. **E.** Compared to the whole genome (wg), stable CHH epimution sites are enriched for CWA trinuleotides, which is a prefer substrate for CMT2 binding.

#### Spontaneous epimutations accumulate neutrally over generations in all cytosine contexts

We started by plotting genome-wide (global) 5mC divergence (*D*) against divergence time (Δt). *D* increases as a function of Δt in all pedigrees (**Fig. 3D**). The increase is rapid for context CG but appears to be low, or even absent, in contexts CHG and CHH. Similar observations have previously led to the hypothesis that the transgenerational heritability of spontaneous epimutations may be restricted to CG dinucleotides [13] [11], perhaps as a consequence of the preferential reinforcement of CHG and CHH methylation during sexual reproduction [28] [29]. Using heuristic arguments it had further been suggested that CG epimutations accumulate neutrally, at least at the genome-wide scale; meaning that 5mC gains and loss in this context are under no selective constraints [13]. However, these hypotheses have never been tested formally due to a lack of analytical tools.

To address this, we used *AlphaBeta* to fit four competing models (ABneutral, ABmm, ABuu, ABnull) to the diverence data of each pedigree (**Fig. 3C**). As mentioned above, model ABneutral assumes that spontaneous 5mC gains and losses accumulate neutrally across generations, ABmm assumes that the accumulation is partly shaped by selection against spontaneous losses of 5mC, ABuu assumes that the accumulation is partly shaped selection against spontaneous gains, and ABnull is the null model of no accumulation (see Methods).

Model comparisons revealed that ABneutral provides the best fit to the 5mC divergence data in context CG in all pedigrees (**Fig. 3C, Tables S2-S4**). This was true at the genome-wide scale (global) as well as at the sub-genomic scale (exons, promoters, TEs). Globally, ABneutral explained between 77% and 90% of the total variance in *D*, indicating that a neutral epimutation model provides a good and sufficient description of the molecular process that generates heritable 5mC changes over time. Interestingly, we also detected, for the first time, highly significant accumulation of neutral epimutations in contexts CHG and CHH (**Fig. 3C, Tables S2-S4**). However, the detection of these accumulation patterns was mainly restricted to MA1_1, the largest of the three pedigrees in terms of both sample size (*N* =26) and divergence times (max. Δt=64), and to some extent also to MA3, the second largest of the three pedigrees (*N* = 13, max. Δt=22). Hence, sufficiently powered studies appear to be necessary to be able detect the accumulation of heritable epimutations in non-CG contexts.

The observation that CHH epimutations are cummulative over generations is some-what suprising, given that CHH methylation is mainly targeted by the *de novo* RNA directed DNA methylation pathway (RdDM), which should prevent the formation stable epimutations, particularly those originating from spontaneous losses of methylation. We therefore examined specific CHH sites that show stable methylation status changes over time (**Fig. 3E**), and found overwhelming evidence that these sites do not correspond to RdDM targets (**Fig. 3F**. Instead they are targeted by CMT2, an enzyme that maintains methlyation states at a subset of CHG and CHH sites, independently RdDM (**Fig. 3G-H**. The preferential targeting of these CHH sites by CMT2 provides a molecular explanation why stochastic losses of DNA methylation are maintained, rather than being re-established *de novo* over generation time.

#### Robust estimates of the rate and spectrum of spontaneous epimutations

We examined the estimated epimutation rates corresponding to the best-fitting models from above (**Fig. 4A, Tables S2-S4**). Globally, we found that the CG methylation gain rate (*α*) is 1.4 · 10^−4^ per CG per haploid genome per generation on average (range: 8.6 · 10^−5^ to 1.94 · 10^−4^) and the loss rate (*β*) is 5.7 · 10^−4^ on average (range: 2.5 · 10^−4^ to 8.3·10^−4^). Using data from pedigree MA1_1, we also obtained the first epimutation rate estimates for contexts CHG and CHH. The gain and loss rates for CHG were 3.5 · 10^−6^ and 5.8 · 10^−5^ per CHG per haploid genome per generation, respectively; and for CHH they were 1.9 · 10^−6^ and 1.6 · 10^−4^ per CHH per haploid genome per generation. Hence, transgenerationally heritable CHG and CHH epimutations arise at rates that are about 1 to 2 orders of magnitude lower than CG epimutations in *A. thaliana*, which is reflected in the relatively slow increase of 5mC divergence in non-CG contexts over generation time (**Fig. 3D**).

**Figure 4:**
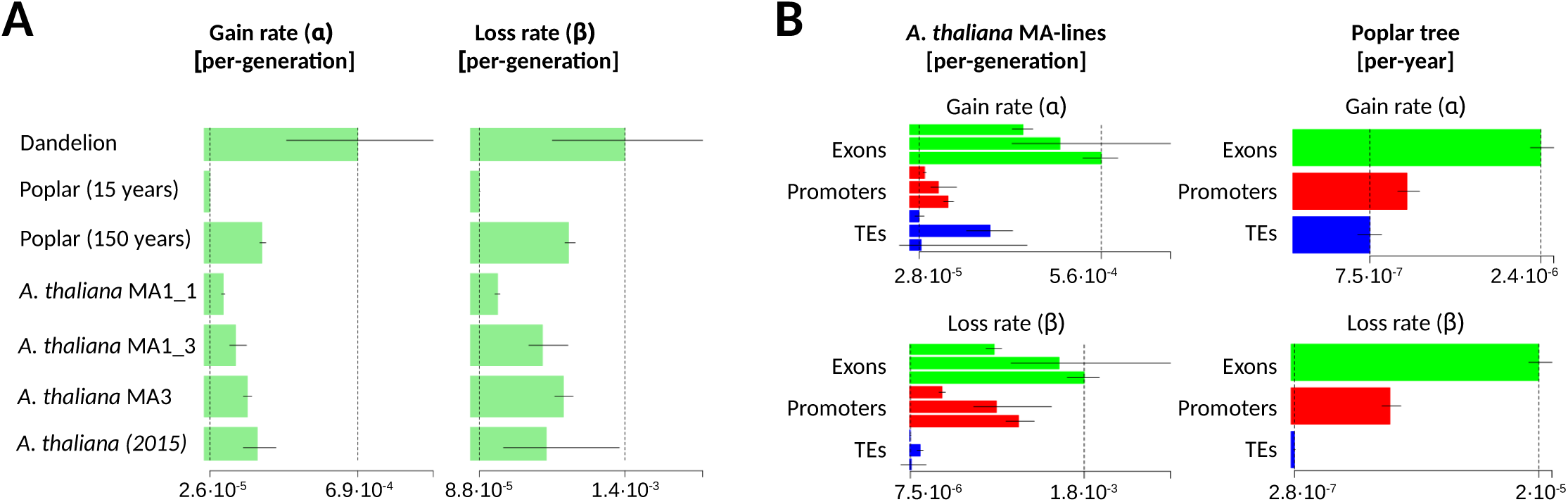
Comparisons of the CG epimutation rates. **A.** Shown are the estimates (+/- 95% confidence intervals) of the genome-wide (global) CG epimutation rates for the different pedigrees. For comparison, we also show the range of previous estimates from *A. thaliana* MA lines (*A. thaliana* (2015)); see van der Graaf et al. (2015). In poplar, the estimated per-year epimutation rates were converted to per-generation rates by assuming a generation time of 15 years and 150 years. The gain and loss rates are all well within one order of magnitude of each other, and differences are mostly within estimation error. The epimutation rate estimates in Dandelion are expected to slightly biased upward due to the application of a diploid model to triploid data (see text). The dashed vertical lines mark off the lower and upper range of the point estimates. **B.** Side-by-side comparison of ‘germline’ and somatic epimutation rate estimates (+/- 95% confidence intervals) in *A. thaliana* MA-lines and poplar, respectively, for selected genomic features. The rank ordering of the magnitude of these rates is similar. For *A. thaliana*, the order of presentation of the pedigrees is MA3, MA1_3 and MA1_1 (from bottom to top within each feature). Feature-specific rates could not be obtained in Dandelion since no annotated assembly is currently available.

In addition to global estimates, we also assessed the gain and loss rates for selected genomic features (exons, promoters, TEs). In line with previous analyses [13], we found striking and consistent rate differences, with exon-specific epimutation rates being 2 to 3 orders of magnitude higher than TE-specific rates (**Fig. 4B, Tables S2-S4**). Interestingly, this trend was not only restricted to CG sites, but was also present in contexts CHG and CHH. This later finding points to yet unknown sequence or chromatin determinants that affect the 5mC fidelity of specific regions across cell divisions, independently of CG, CHG and CHH methylation pathways.

The CG epimutation rates reported here differ slightly from our published estimates [13] (**Fig. 4A Tables S3-S4**). This discrepancy is mainly the result of key differences in the data pre-processing. Application of *AlphaBeta* to published pre-processed samples, yielded similar results to those reported previoulsy, indicating that the statistical inference itself is comparable. Unlike past approaches, we utilized the recent *MethylStar* pipeline (https://github.com/jlab-code/MethylStar) for data pre-processing and methylation state calling. This pipeline yields a substantial increase in the number of high-confidence cytosine methylation calls that can be retained for downstream epimutation analysis **Tables S5**. The benefit of this boost in sample size is reflected in the reduced variation in the *α* and *β* estimates across MA pedigree compared with previous reports [13] (**Fig. 4A, Tables S2-S3**).

### Analysis of somatic epimutations in poplar

Despite the above quantitative insights into the rate and spectrum of spontaneous epimutation in *A. thaliana*, it remains unclear how and where these epimutations actually originate in the plant life cycle. One hypothesis is that they are the result of imperfect 5mC maintenance during the mitotic replication of meristematic cells which give rise to all above and below ground tissues, including the ‘germline’ (**Fig. 1**). As the germline is believed to be derived quite late in development from somatic precursors, somatic epimutations that accumulate during aging can subsequently be passed to offspring. An alternative hypothesis is that heritable epimutations orginate as a by-product of sRNA-mediated reinforcement errors in the sexual cell linages. One way to distinguish these two possibilities is to study epimutational processes in systems that bypass or exclude sexual reproduction.

Long-lived perennials, such as trees, represent a powerful system to explore this. As the tree branching structure can be interpreted as an intra-organismal phylogeny of different somatic cell lineages, it it possible to track mutations and epimutations and their patterns of inheritances across different tree sectors. Recently, there has been a surge of interest in characterizing somatic nucleotide mutations in trees using whole genome sequencing data [30] [31] [32] [33]. These studies have shown that fixed mutations arise sequentially in different tree sectors, thus pointing at a shared meristematic origin.

To facilate the first insights into epimutational processes in long-lived perennials, we applied *AlphaBeta* to MethylC-seq leaf samples (∼ 41.1X coverage) from 8 seperate branches of a single poplar (*Populus trichocarpa*) tree (**Fig. 5**, Methods). The tree features two main stems (here refered to as tree 13 and tree 14), which were originally thought to be two separate trees. However, both stems are stump sprouts off an older tree that was knocked down about 350 years ago. In other words, tree 13 and tree 14 are clones that have independently diverged for a long time. Four branches from each tree were chosen and aged by coring at the points where each branch meets the main stem as well as at the terminal branch (**Fig. 5A-B**, Method). Age-dating of the bottom sector of the tree proved particularly challenging because of heart rot, rendering estimates of the total tree age imprecise. However, an informed guess places the minimum age of the tree at about 250 years.

**Figure 5:**
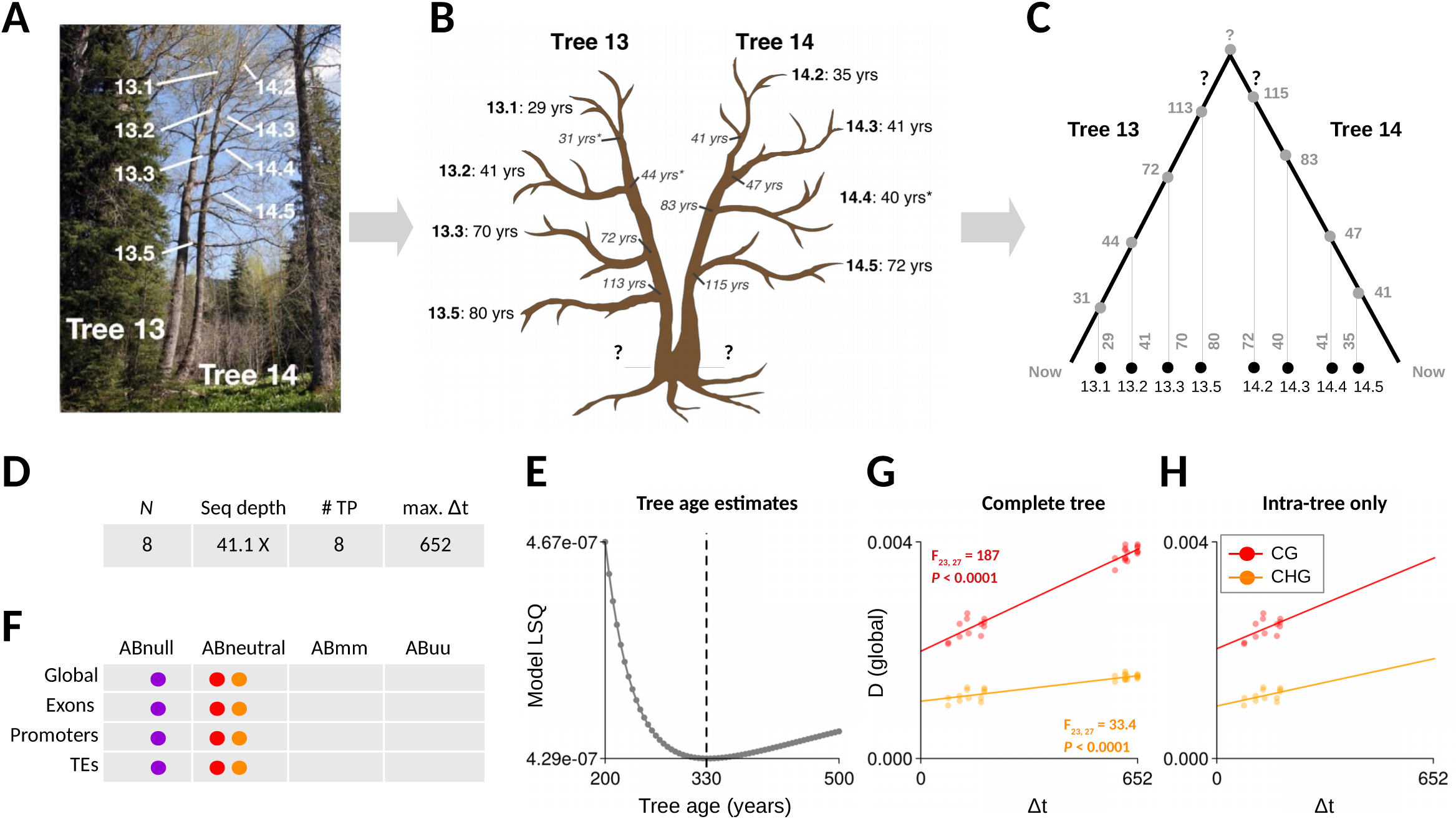
Analysis of somatic epimutations in poplar. **A.** A single poplar (*P. trichocarpa*) tree was analyzed. Originally believed to be two separate trees (tree 13 and tree 14), they are actually part of one tree. Hence tree 13 and tree 14 are clones that have independently diverged. Four branches from each tree were chosen and aged by coring **B.** Schematic representation of tree 13 and tree 14, along with the age estimates obtained from the indicated coring sites. Coring was performed where each of the lateral shoots meets the main stems. Age-coring proved technically challenging at the bottom of the tree and led to unintelligible ring counts. An educated guess places the age of the tree between 250 and 400 years. **C.** The tree can be presented as an intra-organismal phylogeny. The branch point-times and the branch lengths are known from the coring data, with the exception of the bottom sector of the tree (indicated by question marks). Leaf methylomes were collected from each of the selected branches and served as input for *AlphaBeta*. **D.** Overview of the data: *N* is the total number of sequenced samples; Seq depth is the average sequence depths of the samples; # TP is the number of unique time-points that are sampled; max. Δt is the maximum divergence time (in years) between leaf samples. **E.** *AlphaBeta* was fitted to the global CG methylation divergence data of the complete tree data treating the unknown age of the tree as an additional model parameters. Model residual sums of squares (LSQ) were minimized at an age of 330 years, which is our estimate of the age of the tree. **F.** Model comparisons indicate that somatic epimutations accumulate neutrally in context CG (red) and CHG (orange) during aging, both at the global scale as well as within specific genomic features (exons, promoters, TEs). We found no evidence for epimutation accumulation in context CHH (purple). **G-F.** Shown are the fits of model ABneutral to the global CG (red) and CHG (orange) methylation divergence data of the complete tree (intra-tree + inter-tree, **G.**), as well as for tree 13 and tree 14 separately (intra-tree, **H.**).

#### Inferring total tree age from inter-branch leaf methylome data

We used the coring-based age measurements from each of the branches along with the branch points to calculate divergence times (Δt) between all pairs of leaf samples (**Fig. 5C**). We did this by tracing back their ages (in years) along the branches to their most recent common branch point (i.e. ‘founder cells’) (**Fig. 2A, Fig. 5A-C**). The calculation of the divergence times for pairs of leaf samples originating from tree 13 and tree 14 was not possible since the total age of the tree was unknown. To solve this problem, we included the total age of the tree as an additional unknown parameter into our epimutation models. Our model estimates revealed that the total age of the tree is approximately 330 years (**Fig. 5E**), an estimate that fits remarkably well with the hypothesized age window (between 250 and 350 years). Furthermore, the model fits provided overwhelming evidence that somatic epimuations, in poplar, accumulate in a selectively neutral fashion during aging, both at the genome-wide scale (globally) as well as at the sub-genomic scale (exons, promoters, TEs) (**Fig. 5F, Table S1**). This was true for both contexts CG and CHG, which displayed clear 5mC divergence over time (**Fig. 5G**). To rule out that these accumulation patterns are not dominated by our age estimate, we also examined the accumulation patterns within tree 13 and tree 14 separately. We found similar accumulation slopes as well as epimuation rate estimates (**Fig. 5H**).

Taken together, our results provide the first evidence that neutral epimutations accumulate during the somatic development of trees, and that inter-branch leaf methylome data, in conjuncion with our statistical models, can be used as a molecular clock to age-date trees. With sufficiently large sample sizes it should be feasible to infer and date the complete tree branching structure, possibly even without any coring data. Such efforts are currently underway.

#### Estimates of the rate and spectrum of somatic epimutations

We examined the somatic epimutation rate estimates from our best fitting model from the complete tree analysis. At the genome-wide scale, we found that the 5mC gain and loss rates in context CG are 1.7·10^−6^ and 5.8·10^−6^ per site per haploid genome per year, repectively; and 3.3·10^−7^ and 4.1·10^−6^ in context CHG. Interestingly, these *per year* CG epimutation rates are only about two orders of magnitude lower than the *per generation* rates in *A. thaliana* MA-lines. Assuming an average generation time of about 15 years to 150 years in poplar [34], its expected per-generation CG epimutation rate would be between ∼ 10^−5^ to ∼ 10^−4^, which is within the same order of magnitude to that of *A. thaliana* (∼ 10^−4^) (**Fig. 4A**). This close similarity is remarkable given that poplar is about ∼100 times larger and its life-cycle ∼1000 times longer than that of *A. thaliana*. Similar insights were reached in a recent comparison of the per-generation nucleotide mutation rates between Oak (*Quercus rubur*) and *A. thaliana* [32], which were also found to be remarkably close to each other. Taken together, these findings support the emerging hypothesis that meristematic cells must undergo very few divisions during tree aging, and that they are shielded from the century-long exposure to enviornmentnal mutagens, such as UV radiation [33].

To assess whether the accumulation dynamics of somatic epimutations in poplar differs between genomic features, we examined in more detail the estimated rates and spectra for exons, promoters and TEs (**Fig. 4B**). Focusing on context CG, we found considerable rate differences. The gain rates for exons, promoters and TEs were 2.4·10^−6^, 1.1·10^−6^, and 7.5·10^−7^ per site per haploid genome per year, respectively, and the loss rates were 2 · 10^−5^, 8 · 10^−6^, and 2.8 · 10^−7^. Intriguingly, the rank order of these rates was similar to what we had observed for germline epimutations in *A. thaliana*, with exons showing the highest combined rates, followed by promoters and then TEs (**Fig. 4B**). These findings suggest that the fidelity of feature-specific DNA methylation maintenance is deeply conserved across angiosperms, and that feature-specific epimutation accumulation patterns are established during somatic development, rather than being a byproduct of selective reinforcement of DNA methylation in the germline or early zygote. Identifying *cis*- and *trans*-determinants that affect local epimutation rates seems to be an important next challenge [11].

### Analysis of spontaneous epimutations in asexually-derived dandelion MA-lines

Our analysis of *A. thaliana* and poplar revealed striking similarities in the rate and spectrum of CG epimutations. To faciliate further inter-specific comparisons of epimutational processes among angiosperms, particulary across different mating systems, we generated novel MA-lines in an asexual dandelion (*Taraxacum officinale*) genotype (AS34) [35](**Fig. 6A**). Apomictic dandelions are triploid and produce asexually via clonal seeds in a process that involves unreduced egg cell formation (diplospory), parthenogenic embryo development and autonomous endosperm formation, resulting in genetically identical offspring [36]. Using single-seed descent from a single apomictic triploid founder genotype, 8 replicated lineages were propagated for 6 generations, and 5mC measurements were obtained from each generation (**Fig. 6A**).

**Figure 6:**
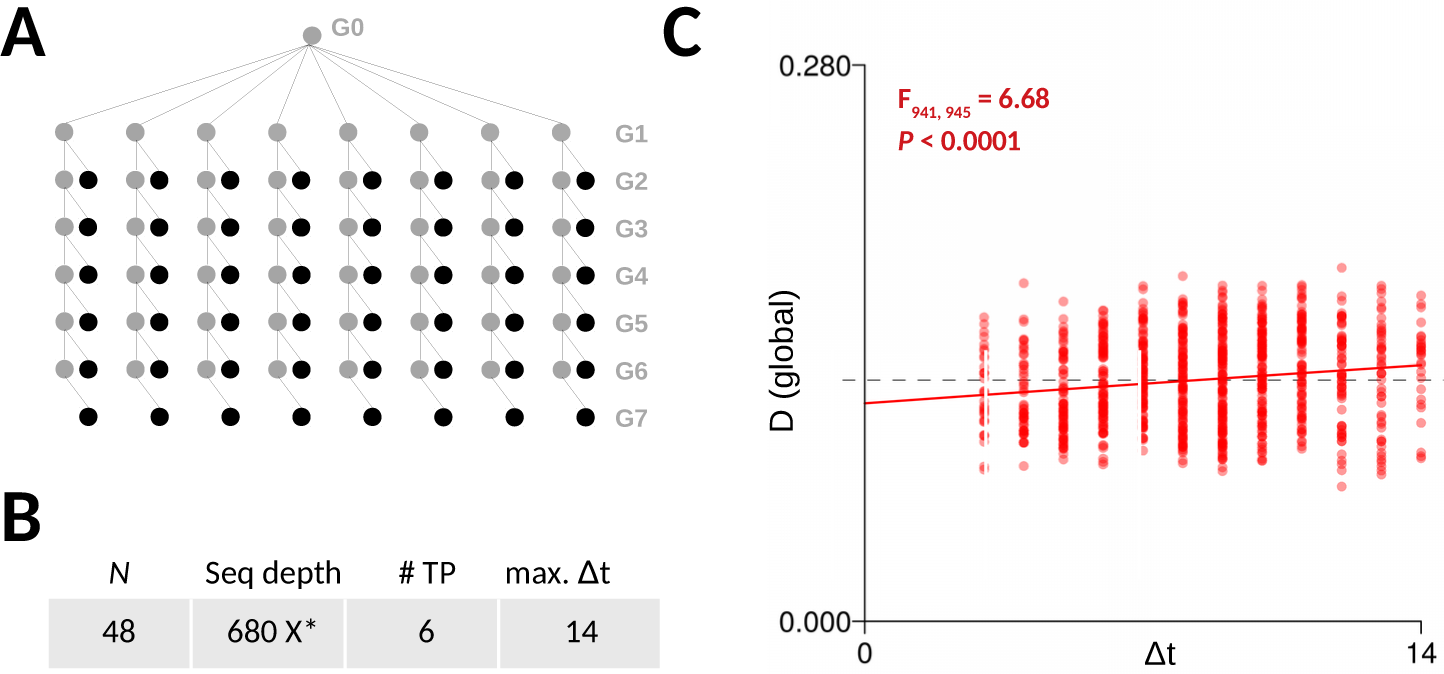
Analysis of GG epimutations in apomictic Dandelion. Using single-seed descent from a single apomictic triploid founder genotype, 8 replicated lineages were propagated for 6 generations. DNA methylation measurements were obtained using epigenotyping-by-sequencing (epiGBS). **B.** Overview of the data: *N* is the total number of sequenced samples; Seq depth is the average sequence depths of the samples; # TP is the number of unique time-points that are sampled; max. Δt is the maximum divergence time (in generations) between samples. *Note: the calculation of average read coverage was based only on interrogated cytosines as epiGBS does not yield any genome-wide data. **C.** Model fits to the CG divergence data. Highly significant increases in 5mC divergence (D) over generation time (Δt) were detected in all sequence contexts, despite the relatively large variation in 5mC divergence patterns (see text).

The total dataset was relatively large, with 48 sequenced samples and a maximum divergence time of 14 generations (**Fig. 6B**). 5mC measurements were obtained using epigenotyping-by-sequencing (epiGBS) [27] (see Methods). Since there is currently no published dandelion reference assembly, local assemblies were generated *de novo* from the epiGBS short reads, and served as basis for cytosine methylation calling [27]. With this approach, ∼24000 measured cytosines were shared between any two sample pairs on average, and were used to calculate pair-wise CG methylation divergence *D*.

Plotting *D* against divergence time (Δt) revealed considerable measurement variation across samples (**Fig. 6C**). This large variation could have several possible sources: First, methylation state calling was based on local assemblies rather than on reference-based alignments. Second, epiheterozygotes in this triploid genotype could not be effectively distinguished on the basis of the observed methylation levels, which introduce uncertainties in the calculation *D*. Third, early implementations of the epiGBS protocol could not distinguish PCR duplicates, a problem that has since been solved [37].

Despite these limitations, application of *AlphaBeta* to the CG divergence data revealed strong statistical evidence for epimutation accumulation over time (*F*_941,945_=6.68, *p* < 0.0001). Consistent with *A. thaliana* and poplar, a neutral epimutation model (AB-neutral) provided the best fit to the data. Based on these model fits, we examined CG epimutation rate estimates. The global gain rate and loss rates were 6.9 · 10^−4^ and 1.4 · 10^−3^ per CG site per haploid genome per generation (**Fig. 4**). However, these estimates should be taken with caution since we applied *AlphaBeta*’s diploid models to data from a triploid species. Although this model mis-specification should have little impact in a clonal system, the resulting ‘per-haploid’ rate estimates are expected to be slightly biased upward.

Keeping this caveat in mind, our results show that the dandelion per-generation CG epimutation rates are close to those obtained in *A. thaliana* and poplar (**Fig. 4A**), and at least within the same order of magnitude. This finding is probably a direct consequence of the fact that DNA methylation maintenance pathways are highly conserved across angiosperms [5] [38]. Moreover, it lends further support to the hypothesis that sexual reproduction has no major impact on the formation and inheritance of spontaneous epimutations. To test this hypthesis in more detail, experimental approaches should be taken that study epimutational processes in a single species/genotype, which is propagated both sexual and asexually over multiple generations. Such a set up would permit a more direct comparison.

## Discussion

Accurate estimates of the rate and spectrum of spontaneous epimuations are essential for understanding how DNA methylation diversity arises in the context of plant evolution, development and aging. Here we presented *AlphaBeta*, a computational method for obtaining such estimates from pedigree-based high-throughput DNA methylation data. Our method requires that the topology of the pedigree is known. This requirement is typically met in the experimental construction of mutation accumulation lines (MA-lines) that are derived through sexual or clonal reproduction. However, we demonstrated that *AlphaBeta* can also be used to study somatic epimutations in long-lived perennials, such as trees, using leaf methylomes and coring data as input. In this case, our method treats the tree branching structure as an intra-organismal phylogeny of somatic lineages and uses information about the epimutational history of each branch.

To demonstrate the versatility of our method, we applied *AlphaBeta* to very diverse plant systems, including multi-generational DNA methylation data from selfing- and asexually derived MA-lines of *A. thaliana* and dandelion, as well as intra-generational DNA methylation data of a poplar tree. Our analysis led to several novel insights about epimutational processes in plants. One of the most striking findings was the close similarity in the epimutation landscapes between these very different systems. Close similarities were observed in the per-generation CG epimutation rates between *A. thaliana*, dandelion and poplar both at the genome-wide as well as at the subgenomic scale. Any detected rate differences between these different systems were all within one order of a magnitude of each other, and as such practically indistinguishable from experimental sources of variation. As a reference, variation in epimutation rate estimates across different *A. thaliana* mutation accumulation experiments vary up to 75% of an order of a magnitude. Clearly, larger sample sizes are needed along with controlled experimentally comparisons to be able to identify potential biological causes underlying subtle epimutation rate differences between species, mating systems, genotypes or environmental treatments. Furthermore, the close similarity between sexual and asexual (or somatic) systems reported here provide indirect evidence that transgenerationally heritable epimutations originate mainly during mitotic rather than during meiotic cell divisions in plants.

Our application of *AlphaBeta* to poplar also provided the first proof-of-principle demonstration that leaf methylome data, in combination with our statistical models, can be employed as a molecular clock to age-date trees or sectors of trees. Anaytically, this is similar to infering the branch lengths of the underlying pedigree (or phylogeny). With sufficiently large sample sizes it should be possible to achieve this with relatively high accuracy, and extent this inference to the entire tree structure. The comparatively high rate of somatic and germline-epimutations are instrumental in this as they provide increased temporal resolution over classical DNA sequence approaches, which rely on rare *de novo* nucleotide mutations. Our methodological approach should be applicable, more generally, to any perennial or long-lived species. We are currently extending the *AlphaBeta* tool set to faciliate such analyses.

Analytically, *AlphaBeta* is not restricted to the analysis of plant data. The method could also be used to study epimutational processes in tumor clones based on animal single cell WGBS data. Such datasets are rapidly emerging [39]. In this context, *AlphaBeta* could be instrumental in the inference of clonal phylogenies and help calibrate them temporally. Such efforts may complement current pseudotemporal ordering (or trajectory inference) methods and lineage tracing strategies in single-cell methylation data [40] [41].

The implementation of *AlphaBeta* is relatively straight-forward. The starting point of the method are methylation state calls for each cytosine. These can be obtained from any methylation calling pipeline. In the data applications presented here we used *AlphaBeta* in conjunction with *MethylStar* (https://github.com/jlab-code/MethylStar) (**Figure2**), which is an efficient pipeline for the analysis of population-scale WGBS data and features a HMM-based methylation state caller [42]. Application of this pipeline leads to up a substantial increase in the number of high-confidence cytosines methylation calls for epimutation rate inference compared with more convential methods. We therefore recommend using *AlphaBeta* in conjunction with *MethylStar*. Software implementing *AlphaBeta* is available as a Bioconductor R package at http://bioconductor.org/packages/3.10/bioc/html/AlphaBeta.html

## Methods

### A. thaliana MA-lines data

#### Plant material

For MA3, seeds were planted and grown in 16-hour day lengths and samples were harvested from young above ground tissue. Tissue was flash frozen in liquid nitrogen and DNA was isolated using a Qiagen Plant DNeasy kit (Qiagen, Valencia, CA, USA) according to the manufacturer’s instructions. For MA1_1 and MA1_3, a detailed description of growth conditions and plant material can be found in the original publications [17] [13].

#### Sequencing and data processing

For MA3, MethylC-seq libraries were prepared according to the protocol described in Urich et al. [43]. Libraries were sequenced to 150-bp per read at the Georgia Genomics & Bioinformatics Core (GGBC) on a NextSeq500 platform (Illumina). Average sequencing depth was 20.8X among samples (**Table S1**). For MA1_1 and MA1_3, FASTQ files (*.fastq) were downloaded from (URL). All data processing and methylation state calling was performed using the *MethylStar* pipeline (github.com/jlab-code/MethylStar). Summary statistic for each sample can be found in **Table S1**.

### Poplar data

#### Tree coring

The tree used in this study was located at Hood River Ranger District [Horse Thief Meadows area], Mt. Hood National Forest, 0.6 mi south of Nottingham Campground off OR-35 at unmarked parking area, 500′ west of East Fork Trail nbr. 650 across river, ca. 45.355313, -121.574284.Tree cores were originally collected from the main stem and five branches in April 2015 at breast height (∼ 1.5 m) for standing tree age using a stainless-steel increment borer (5 mm in diameter and up to 28 cm in length). Cores were mounted on grooved wood trim, dried at room temperature, sanded and stained with 1% phloroglucinol following the manufacturer’s instructions (https://www.forestry-suppliers.com/Documents/1568_msds.pdf).

Annual growth rings were counted to estimate age. For cores for which accurate estimates could not be made from the 2015 collection, additional collections were made in spring 2016. However, due to difficulty in collecting by climbing, many of the cores did not reach the center of the stem or branches (pith) and/or the samples displayed heart rot. Combined with the difficulty in demarcating rings in porous woods such as poplar Populus (cite 40,41), accurate measures of tree age or branch age were challenging.

#### Sequencing and data processing

A single MethylC-seq library was created for each branch from leaf tissue. Libraries were prepared according to the protocol described in Urich et al [43]. Libraries were sequenced to 150-bp per read at the Georgia Genomics & Bioinformatics Core (GGBC) on a NextSeq500 platform (Illumina). Average sequencing depth was 41.1x among samples. MethylC-seq reads were aligned using Methylpy v1.3.2 [44]. Alignment was to the new Stettler14 assembly of *P. trichocarpa*, as described in (see companion paper). Starting from the BAM files (*.bam) the *MethylStar* pipeline was used for further data processing and methylation state calling.

### Dandelion MA-lines data

#### Plant material

Starting from a single founder individual, eight replicate lineages of the apomictic common dandelion (*Taraxacum officinale*) genotype AS34 [35] were grown for six generations via single-seed descent under common greenhouse conditions. Apomictic dandelions are triploid and produce asexually via clonal seeds in a process that involves unreduced egg cell formation (diplospory), parthenogenic embryo development and autonomous endosperm formation, resulting in genetically identical offspring [36]. Seeds were collected from each of the 48 plants in the six-generation experiment and stored under controlled conditions (15 degrees Celcius and 30% RH). After the 6th generation, from each plant in the pedigree a single offspring individual was grown in a fully randomized experiment under common greenhouse conditions. Leaf tissue from a standardized leaf was collected after five weeks, flash frozen in liquid nitrogen and stored at -80 degrees Celsius until processing.

#### Sequencing and data processing

DNA was isolated using the Macherey-Nagel Nucleospin Plant II kit (cell lysis buffer PL1). DNA was digested with the PstI restriction enzyme and epiGBS sequencing libraries were prepared as described elsewhere [27]. Based on genotyping-by-sequencing [45], epiGBS is a multiplex Reduced-Representation Bisulphite Sequencing (RRBS) approach with an analysis pipeline that allows for local reference construction from bisulphite reads, which makes the method applicable to species for which a reference genome is lacking [27]. PstI is a commonly used restriction enzyme for genotyping-by-sequencing, however, its activity is sensitive to CHG methylation in CTGCAG recognition sequence. This makes the enzyme better at unbiased quantification of CG methylation than of CHG methylation [27]. After quantification of the sequencing libraries using a multiplexed Illumina MiSeq Nano run, samples were re-pooled to achieve equal representation in subsequent epiGBS library sequencing. The experimental samples were sequenced on two Illumina HiSeq 2500 lanes (125 cycles paired-end) as part of a larger epiGBS experiment which consisted of a total of 178 samples that were randomized over the two lanes. Because of inadequate germination or due low sequencing output (library failure), four of the 48 samples were not included in the downstream analysis.

#### DNA methylation analysis

Sequencing reads were demultiplexed (based on custom barcodes) and mapped against a dandelion pseudo-reference sequence that was generated *de novo* from PstI-based epiGBS [27]. This pseudo-reference contains the local reference of PstI-based epiGBS fragments as inferred from the bisulphite reads. Methylation variant calling was based on SAMtools mpileup and custom python scripts, following a similar approach as described in van Gurp et al. [27]. For downstream analysis, we included only those cytosines that were called in at least 80% of the samples. In addition, cytosine positions that did not pass the filtering criteria for all generations were removed.

To obtain methylation status calls, we implemented a one-tail binomial test as previously described [13]. Multiple testing correction was performed using the Benjamini-Yekutiely method, and the false discovery rate (FDR) was controlled at 0.05. All statistical tests for obtaining methylation status calls of the samples were conducted within the SciPy ecosystem.

### The AlphaBeta method

#### Calculating 5mC divergence

For the *i*th sequenced sample in the pedigree, let *s*_*ik*_ be the observed methylation state at the *k*th locus (*k* = 1 … *N*). Here, the *N* loci can be individual cytosines or pre-defined regions (i.e. cluster of cytosines). We assume that *s*_*ik*_ takes values 1, 0.5 or 0, according to whether the diploid epigenotype at that locus is *m/m, m/u, u/u*, respectively, where *m* is a methlylated and *u* is an unmethylated epiallele. Using this coding, we calculate the total 5mC divergence, *D*, between any two samples *i* and *j* in the pedigree as follows:

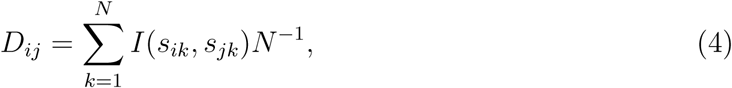

where *I*(·) is an indicator function, such that

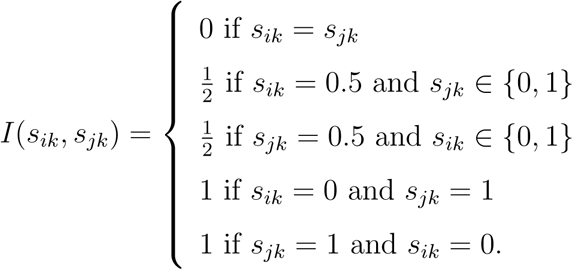

#### Modelling 5mC divergence

We model the 5mC divergence as

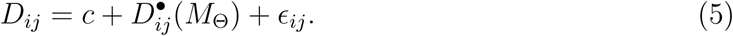

Here *ϵ*_*ij*_ ∼ *N* (0, *σ*^2^) is the normally distributed residual error, *c* is the intercept, and 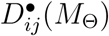 is the expected divergence between samples *i* and *j* as a function of an underlying epimutation model *M* (·) with parameter vector Θ (see below). We have that

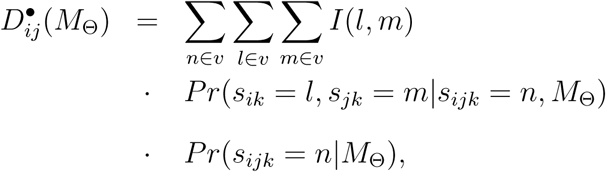

where *s*_*ijk*_ is the methylation state at the *k* locus of the most recent common ancestor of samples *i* and *j* (**Fig. 3**), and *v* = {0, 0.5, 1}. Since samples *s*_*i*_ and *s*_*j*_ are conditionally independent, we can further write:

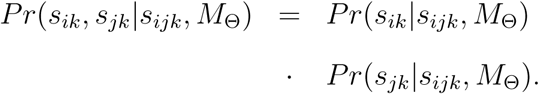

To be able to evaluate these conditional probabilities it is necessary to posit an explicit form for the epimutational model, *M*_Θ_. To motivate this, we define **G** to be a 3 × 3 transition matrix, which summarizes the probability of transitioning from epigenotype *l* to *m* in the time interval [*t, t* + 1]:

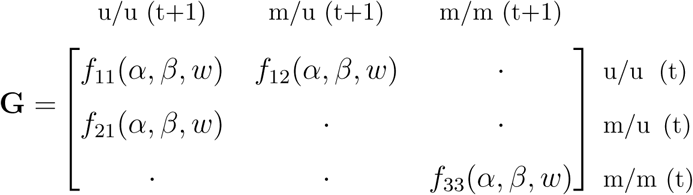

The elements of this matrix are a function of gain rate *α* (i.e. the probability of a stochastic epiallelic switch from an unmethylated to a methylated state within interval [*t, t* + 1]), the loss rate *β* (i.e. the probability of a stochastic epiallelic switch from a methylated to an unmethylated state), and the selection coefficient *w* (*w* ∈ [0, 1]). It can be shown that for a diploid system propagated by selfing, **G** has the form

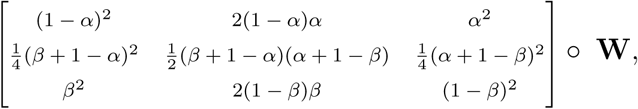

and for systems that are propagated clonally or somatically **G** is:

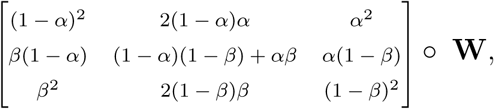

where ∘ is the Hadamard product and **W** is a matrix of selection coefficients of the form

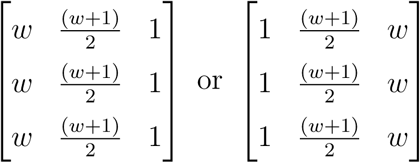

depending on whether selection is against epigenotype *u/u* or *m/m*, respectively. For instance, in the case of selection against epigenotypes *u/u*, the fitness of epihomozygote *u/u* and epiheterozygote *m/u* are reduced by a factor of *w* and (*w* + 1)*/*2, respectively. We incorperate this fitness loss directly into the transition matrix by weighing the transition probabilities to these epigenotypes accordingly. Similar arguments hold for the case where selection is against *m/m*. In the special situation where *w* = 1 we have a neutral model, where epigenotype transitions from time *t* to *t* + 1 are only governed by the rates *α* and *β*, and - in the case of selfing - also by the Mendelian segregation of epialleles *u* and *m*. To ensure that the rows of **G** (i.e. the transition probabilities) still sum to unity in the presence of selection, we redefine **G** using the normalization:

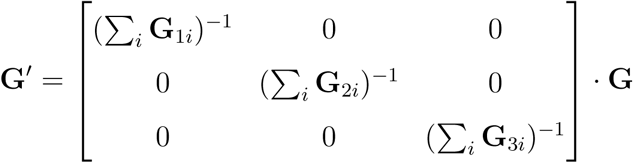

Based on Markov chain theory, the conditional probability *Pr*(*s*_*ik*_|*s*_*ijk*_, *M*_Θ_) can then be expressed in terms of **G**′ as follows:

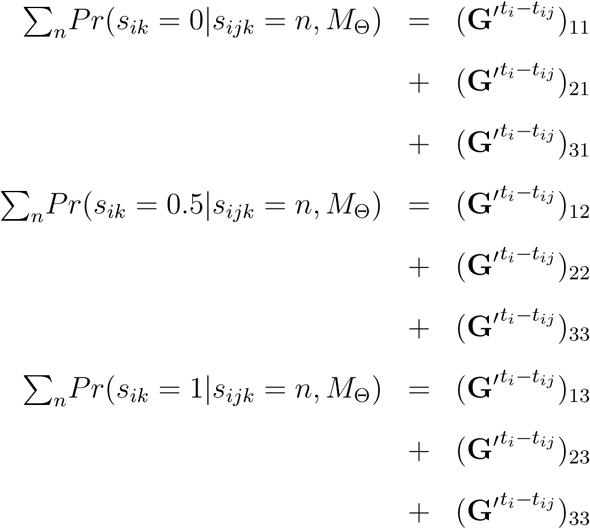

where *t*_*i*_ is the time-point corresponding to sample *i* and *t*_*ij*_ is the timepoint of the most recent common ancestor shared between samples *i* and *j*, (*t*_*ij*_ ≤ *t*_*i*_, *t*_*j*_). Expressions for *Pr*(*s*_*jk*_|*s*_*ijk*_, *M*_Θ_, *t*_*j*_) can be derived accordingly, by simply replacing *t*_*i*_ by *t*_*j*_ in the above equation. Note that the calculation of these conditional probabilities requires repeated matrix multiplication. However, a direct evaluation of these equations is also possible using the fact that

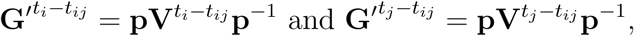

where **p** is the eigenvector of matrix **G**′ and **V** is a diagonal matrix of eigenvalues. For selfing and clonal/somatic systems, these eigenvalues and eigenvectors can be be obtained analytically.

Finally, to derive 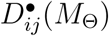, we also need to supply *Pr*(*s*_*ijk*_ = *n*|*M*_Θ_); that is, the probability that any given locus *k* in the most recent common ancestor of samples *i* and *j* is in state *n* (*n* ∈ {0, 0.5, 1}). To do this, consider the methylome of the pedigree founder at time *t* = 1, and let *π* = [*p*_1_ *p*_2_ *p*_3_] be a row vector of probabilities corresponding to states *u/u, u/m* and *m/m*, respectively. Using Markov Chain theory we have

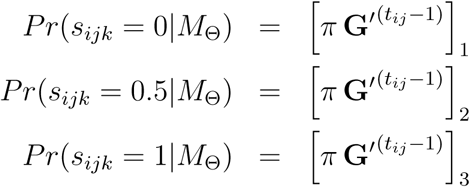

In many situations the most recent common ancestor happens to be the pedigree founder itself, so that *t*_*ij*_ = 1. In the case where the methylome of the pedigree founder has been measured, the probabilities *p*_1_, *p*_2_ and *p*_3_ can be estimated directly from the data using *x*_1_*N* ^−1^, *x*_2_*N* ^−1^ and *x*_3_*N* ^−1^, respectively. Here *x*_1_, *x*_2_ and *x*_3_ are number of loci that are observed to be in states *u/u, u/m, m/m*, and *N* is the total number of loci. Typically, however, *x*_2_ is unknown as most DMP and DMR callers do not output epiheterozygous states (i.e. intermediate methylation calls). Instead, we therefore use

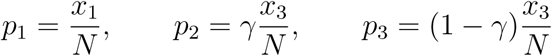

where *γ* ∈ [0, 1] is an unknown parameter. In other words, *AlphaBeta* estimates the proportion of epiheterozygote loci in the pedigree founder during model fitting. An alternative strategy is to obtain *p*_1_, *p*_2_ and *p*_3_ from sample *s*_*ij*_ directly, provided such measurements are available.

#### Model inference

To obtain estimates for Θ, we seek to minimize

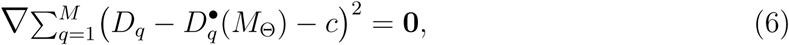

where the summation is over all *M* unique pairs of sequenced samples in the pedigree. Minimization is performed using the “Nelder-Mead” algorithm as part of the optimx package in R. However, from our experience convergence is not always stable, probably because the function 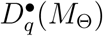 is complex and highly non-linear. We therefore include the following minimization constraint:

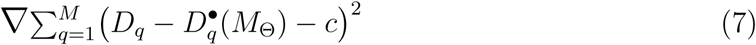

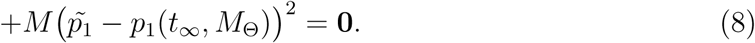

Here *p*_1_(*t*_∞_, *M*_Θ_) is the equilibrium proportion of *u/u* loci in the genome as *t* → ∞. For a selfing system with *w* = 1 we have that

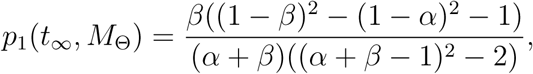

and for a clonal/somatic system it is:

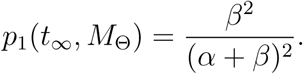

For the case where 0 < *w* ≤ 1 the equations are more complex and are omitted here. Note that the value 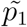 is an empirical guess at these equilibrium proportions. For samples whose methylomes can be assumed to be at equilibrium we have that *p*_1_(*t* = 1) = *p*_1_(*t* = 2) = … = *p*1(*t*_∞_), meaning that the proportion of loci in the genome that are in state *u/u* are (dynamically) stable for any time *t*. Under this assumption, 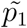 can be replaced by 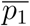, which is the average proportion of *u/u* loci calculated from all pedigree samples.

#### Confidence intervals

We obtain confidence intervals for the estimated model parameters by boostrapping the model residuals. The procedure has the following steps: 1. For the *q*th sample pair *q* (*q* = 1, …, *M*) we define a new response variable 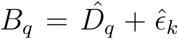, where 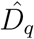 is the fitted divergence for the *q*th pair, and 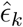 is drawn at random and with replacement from the 1 × *M* vector of fitted model residuals; 2. Refit the model using the new response variable, and obtain estimates for the model parameters. 3. Repeat steps 1. to 2. a large number of times to obtain a boostrap distribution. 4. Use the bootrap distribution from 3. to obtain empirical confidence intervals.

#### Testing for selection

To assess whether a selection model provides a significantly better fit to the data compared to a neutral model, we define

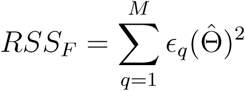

and

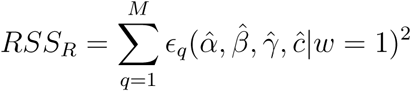

to be the estimated residual sums of squares of the full model and reduced (i.e. neutral) model, respectively, with corresponding degrees of freedom *df*_*F*_ and *df*_*R*_. To test for selection we evaluate the following *F* -statistic:

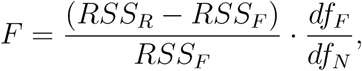

where *df*_*N*_ = *df*_*F*_ − *df*_*R*_. Under the Null *F* ∼ *F* (*df*_*N*_, *df*_*F*_).

## Author’s contributions

FJ and MCT conceptualized the method. FJ, YS and RRH implemented and documented the method. FJ, YS, AS, RRH, JD, TM, BTH, TvG analyzed the data. KV, GT and RJS contributed materials. FJ wrote the paper with input from all coauthors.

## Acknowledgements

We thank Kay Schneitz for discussing plant development with us; Cristina Cipriani for early tests of the optimX package; Keith Slotkin for the sRNA data. FJ and RJS acknowledge support from the Technical University of Munich-Institute for Advanced Study funded by the German Excellent Initiative and the European Seventh Framework Programme under grant agreement no. 291763. FJ is also supported by the SFB/Sonderforschungsbereich924 of the Deutsche Forschungsgemeinschaft (DFG). RJS acknowledges the support from the National Science Foundation (IOS-1546867). RJS is a Pew Scholar in the Biomedical Sciences, supported by the Pew Charitable Trusts.

---

**Table S1:**
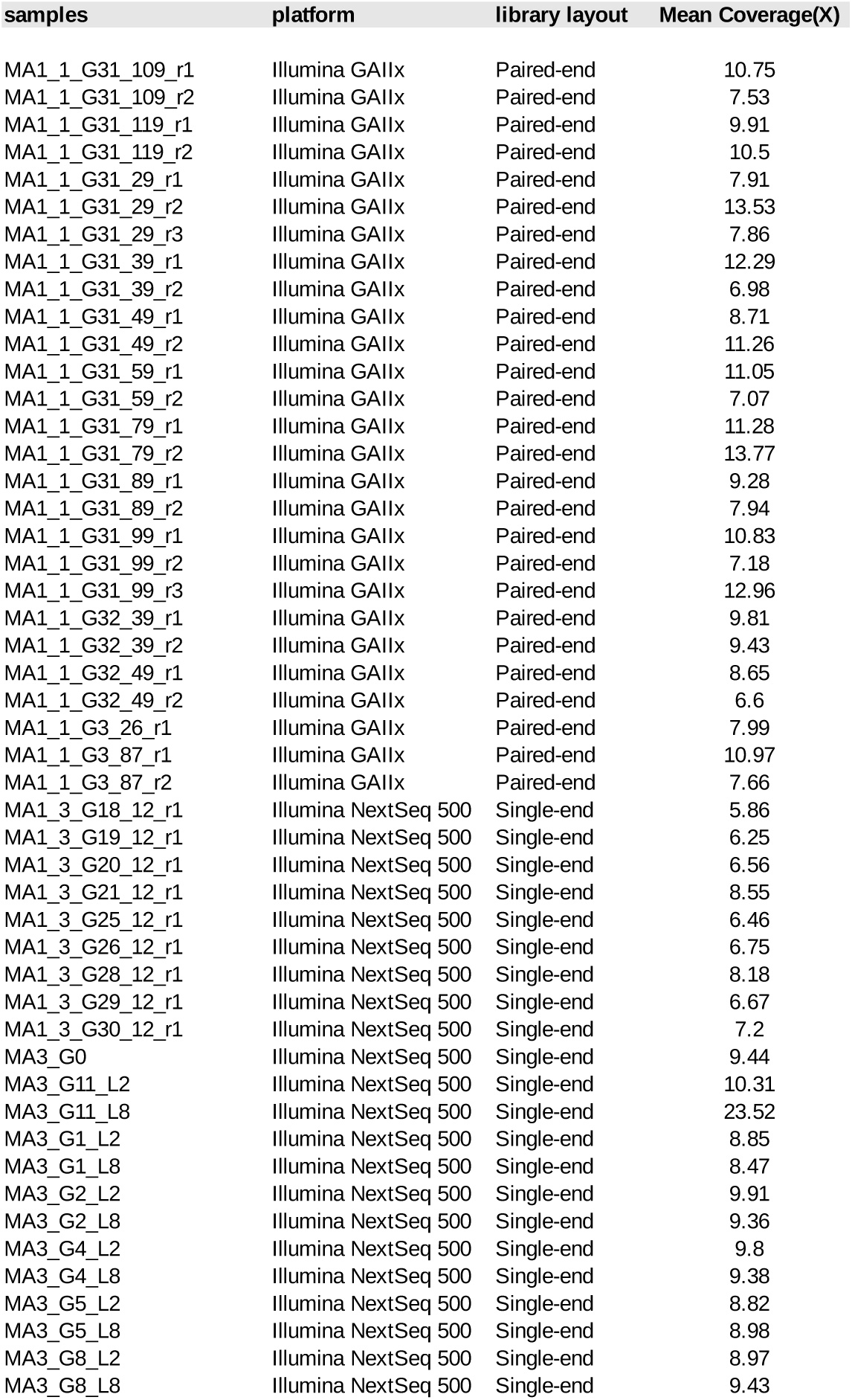
WGBS information for MA pedigrees MA1_1, MA1_3 and MA3.

| **samples** | **platform** | **library layout** | **Mean Coverage(X)** |
| --- | --- | --- | --- |
| MA1_1_G31_109_r1 | Illumina GAIIx | Paired-end | 10.75 |
| MA1_1_G31_109_r2 | Illumina GAIIx | Paired-end | 7.53 |
| MA1_1_G31_119_r1 | Illumina GAIIx | Paired-end | 9.91 |
| MA1_1_G31_119_r2 | Illumina GAIIx | Paired-end | 10.5 |
| MA1_1_G31_29_r1 | Illumina GAIIx | Paired-end | 7.91 |
| MA1_1_G31_29_r2 | Illumina GAIIx | Paired-end | 13.53 |
| MA1_1_G31_29_r3 | Illumina GAIIx | Paired-end | 7.86 |
| MA1_1_G31_39_r1 | Illumina GAIIx | Paired-end | 12.29 |
| MA1_1_G31_39_r2 | Illumina GAIIx | Paired-end | 6.98 |
| MA1_1_G31_49_r1 | Illumina GAIIx | Paired-end | 8.71 |
| MA1_1_G31_49_r2 | Illumina GAIIx | Paired-end | 11.26 |
| MA1_1_G31_59_r1 | Illumina GAIIx | Paired-end | 11.05 |
| MA1_1_G31_59_r2 | Illumina GAIIx | Paired-end | 7.07 |
| MA1_1_G31_79_r1 | Illumina GAIIx | Paired-end | 11.28 |
| MA1_1_G31_79_r2 | Illumina GAIIx | Paired-end | 13.77 |
| MA1_1_G31_89_r1 | Illumina GAIIx | Paired-end | 9.28 |
| MA1_1_G31_89_r2 | Illumina GAIIx | Paired-end | 7.94 |
| MA1_1_G31_99_r1 | Illumina GAIIx | Paired-end | 10.83 |
| MA1_1_G31_99_r2 | Illumina GAIIx | Paired-end | 7.18 |
| MA1_1_G31_99_r3 | Illumina GAIIx | Paired-end | 12.96 |
| MA1_1_G32_39_r1 | Illumina GAIIx | Paired-end | 9.81 |
| MA1_1_G32_39_r2 | Illumina GAIIx | Paired-end | 9.43 |
| MA1_1_G32_49_r1 | Illumina GAIIx | Paired-end | 8.65 |
| MA1_1_G32_49_r2 | Illumina GAIIx | Paired-end | 6.6 |
| MA1_1_G3_26_r1 | Illumina GAIIx | Paired-end | 7.99 |
| MA1_1_G3_87_r1 | Illumina GAIIx | Paired-end | 10.97 |
| MA1_1_G3_87_r2 | Illumina GAIIx | Paired-end | 7.66 |
| MA1_3_G18_12_r1 | Illumina NextSeq 500 | Single-end | 5.86 |
| MA1_3_G19_12_r1 | Illumina NextSeq 500 | Single-end | 6.25 |
| MA1_3_G20_12_r1 | Illumina NextSeq 500 | Single-end | 6.56 |
| MA1_3_G21_12_r1 | Illumina NextSeq 500 | Single-end | 8.55 |
| MA1_3_G25_12_r1 | Illumina NextSeq 500 | Single-end | 6.46 |
| MA1_3_G26_12_r1 | Illumina NextSeq 500 | Single-end | 6.75 |
| MA1_3_G28_12_r1 | Illumina NextSeq 500 | Single-end | 8.18 |
| MA1_3_G29_12_r1 | Illumina NextSeq 500 | Single-end | 6.67 |
| MA1_3_G30_12_r1 | Illumina NextSeq 500 | Single-end | 7.2 |
| MA3_G0 | Illumina NextSeq 500 | Single-end | 9.44 |
| MA3_G11_L2 | Illumina NextSeq 500 | Single-end | 10.31 |
| MA3_G11_L8 | Illumina NextSeq 500 | Single-end | 23.52 |
| MA3_G1_L2 | Illumina NextSeq 500 | Single-end | 8.85 |
| MA3_G1_L8 | Illumina NextSeq 500 | Single-end | 8.47 |
| MA3_G2_L2 | Illumina NextSeq 500 | Single-end | 9.91 |
| MA3_G2_L8 | Illumina NextSeq 500 | Single-end | 9.36 |
| MA3_G4_L2 | Illumina NextSeq 500 | Single-end | 9.8 |
| MA3_G4_L8 | Illumina NextSeq 500 | Single-end | 9.38 |
| MA3_G5_L2 | Illumina NextSeq 500 | Single-end | 8.82 |
| MA3_G5_L8 | Illumina NextSeq 500 | Single-end | 8.98 |
| MA3_G8_L2 | Illumina NextSeq 500 | Single-end | 8.97 |
| MA3_G8_L8 | Illumina NextSeq 500 | Single-end | 9.43 |

**Table S2:** Epimutation rate estimates and model selection results for pedigree MA1_1.

| context | annotation | alpha | beta | beta/alpha | FM | RM | F-value | df RM | df FM | P--value |
| --- | --- | --- | --- | --- | --- | --- | --- | --- | --- | --- |
| CG | global | 8.605897E-05 | 0.0002497981 | 2.903 | ABneutral | Abnull | 461.7119 | 350 | 346 | 2.70488E-137 |
| CG | exon | 0.0003329146 | 0.0008854249 | 2.660 | ABneutral | Abnull | 506.9881 | 350 | 346 | 2.99272E-143 |
| CG | promoter | 4.390624E-05 | 0.0003396206 | 7.735 | ABneutral | Abnull | 574.9184 | 350 | 346 | 2.19508E-151 |
| CG | TE | 2.777663E-05 | 7.500164E-06 | 0.270 | ABneutral | Abnull | 122.3618 | 350 | 346 | 6.001881E-65 |
| CG | global |  |  |  | ABselectUU | Abneutral | 0.4806 | 347 | 346 | 0.4885955 |
| CG | exon |  |  |  | ABselectUU | Abneutral | 0.0453 | 347 | 346 | 0.8316087 |
| CG | promoter |  |  |  | ABselectUU | Abneutral | 1.8765 | 347 | 346 | 0.1716251 |
| CG | TE |  |  |  | ABselectUU | Abneutral | 0.7210 | 347 | 346 | 0.3963924 |
| CG | global |  |  |  | ABselectMM | Abneutral | 0.4731 | 347 | 346 | 0.4920452 |
| CG | exon |  |  |  | ABselectMM | Abneutral | 0.1176 | 347 | 346 | 0.7318225 |
| CG | promoter |  |  |  | ABselectMM | Abneutral | 0.4049 | 347 | 346 | 0.5249724 |
| CG | TE |  |  |  | ABselectMM | Abneutral | 0.1476 | 347 | 346 | 0.7011182 |
| CHG | global | 3.533486E-06 | 5.84628E-05 | 16.545 | ABneutral | Abnull | 16.4449 | 350 | 346 | 2.399368E-12 |
| CHG | exon | 1.537786E-06 | 0.0002547714 | 165.674 | ABneutral | Abnull | 20.6207 | 350 | 346 | 2.958294E-15 |
| CHG | promoter | 2.408792E-06 | 0.0001227248 | 50.949 | ABneutral | Abnull | 19.3769 | 350 | 346 | 2.124209E-14 |
| CHG | TE | 2.124037E-05 | 3.025496E-05 | 1.424 | ABneutral | Abnull | 9.4822 | 350 | 346 | 2.768493E-07 |
| CHG | global |  |  |  | ABselectUU | Abneutral | 0.0000 | 347 | 346 | 1 |
| CHG | exon |  |  |  | ABselectUU | Abneutral | 0.0164 | 347 | 346 | 0.8980838 |
| CHG | promoter |  |  |  | ABselectUU | Abneutral | 0.0000 | 347 | 346 | 1 |
| CHG | TE |  |  |  | ABselectUU | Abneutral | 0.0000 | 347 | 346 | 1 |
| CHG | global |  |  |  | ABselectMM | Abneutral | 0.0000 | 347 | 346 | 1 |
| CHG | exon |  |  |  | ABselectMM | Abneutral | 0.0228 | 347 | 346 | 8.80E-01 |
| CHG | promoter |  |  |  | ABselectMM | Abneutral | 0.0000 | 347 | 346 | 1 |
| CHG | TE |  |  |  | ABselectMM | Abneutral | 0.0000 | 347 | 346 | 1 |
| CHH | global | 1.905105E-06 | 0.0001614412 | 84.741 | ABneutral | Abnull | 11.6186 | 350 | 346 | 7.294017E-09 |
| CHH | exon | 1.09903E-06 | 0.0006017335 | 547.513 | ABneutral | Abnull | 18.1246 | 350 | 346 | 1.577799E-13 |
| CHH | promoter | 1.349004E-06 | 0.0002622809 | 194.426 | ABneutral | Abnull | 19.3769 | 350 | 346 | 2.124209E-14 |
| CHH | TE | 5.54414E-06 | 6.181445E-05 | 11.150 | ABneutral | Abnull | 9.4822 | 350 | 346 | 2.768493E-07 |
| CHH | global |  |  |  | ABselectUU | Abneutral | 0.9755 | 347 | 346 | 0.3240002 |
| CHH | exon |  |  |  | ABselectUU | Abneutral | 0.0000 | 347 | 346 | 1 |
| CHH | promoter |  |  |  | ABselectUU | Abneutral | 0.0000 | 347 | 346 | 1 |
| CHH | TE |  |  |  | ABselectUU | Abneutral | 0.0000 | 347 | 346 | 1 |
| CHH | global |  |  |  | ABselectMM | Abneutral | 0.0000 | 347 | 346 | 1 |
| CHH | exon |  |  |  | ABselectMM | Abneutral | 0.0000 | 347 | 346 | 1 |
| CHH | promoter |  |  |  | ABselectMM | Abneutral | 0.0000 | 347 | 346 | 1 |
| CHH | TE |  |  |  | ABselectMM | Abneutral | 0.0000 | 347 | 346 | 1 |

FM = Full model RM = Reduced model df = degrees of freedom Best performing model

**Table S3:** Epimutation rate estimates and model selection results for pedigree MA1_3.

| context | annotation | alpha | beta | beta/alpha | FM | RM | F-value | df RM | df FM | P--value |
| --- | --- | --- | --- | --- | --- | --- | --- | --- | --- | --- |
| CG | global | 0.0001411325 | 0.0006480785 | 4.592 | ABneutral | Abnull | 26.0964 | 35 | 31 | 1.545386E-09 |
| CG | exon | 0.0004417924 | 0.001566664 | 3.546 | ABneutral | Abnull | 36.7694 | 35 | 31 | 2.352825E-11 |
| CG | promoter | 8.473369E-05 | 0.0009098872 | 10.738 | ABneutral | Abnull | 22.7134 | 35 | 31 | 7.66475E-09 |
| CG | TE | 0.0002361935 | 0.0001108124 | 0.469 | ABneutral | Abnull | 7.9028 | 35 | 31 | 0.0001635841 |
| CG | global |  |  |  | ABselectUU | Abneutral | 0.6312 | 32 | 31 | 0.4329535 |
| CG | exon |  |  |  | ABselectUU | Abneutral | 0.1957 | 32 | 31 | 0.6612462 |
| CG | promoter |  |  |  | ABselectUU | Abneutral | 0.4402 | 32 | 31 | 0.5119528 |
| CG | TE |  |  |  | ABselectUU | Abneutral | 0.0135 | 32 | 31 | 0.9084139 |
| CG | global |  |  |  | ABselectMM | Abneutral | 0.6343 | 32 | 31 | 0.4318425 |
| CG | exon |  |  |  | ABselectMM | Abneutral | 0.2816 | 32 | 31 | 0.5994397 |
| CG | promoter |  |  |  | ABselectMM | Abneutral | 0.2889 | 32 | 31 | 0.5947381 |
| CG | TE |  |  |  | ABselectMM | Abneutral | 0.0163 | 32 | 31 | 0.8993372 |
| CHG | global | NA | NA | NA | ABneutral | Abnull | 0.6354 | 35 | 31 | 0.6410907 |
| CHG | exon | NA | NA | NA | ABneutral | Abnull | 0.4926 | 35 | 31 | 0.7411286 |
| CHG | promoter | NA | NA | NA | ABneutral | Abnull | 0.7356 | 35 | 31 | 0.5747893 |
| CHG | TE | NA | NA | NA | ABneutral | Abnull | 0.3041 | 35 | 31 | 0.8729979 |
| CHG | global |  |  |  | ABselectUU | Abneutral | 0.0271 | 32 | 31 | 0.8703212 |
| CHG | exon |  |  |  | ABselectUU | Abneutral | 0.0000 | 32 | 31 | 1 |
| CHG | promoter |  |  |  | ABselectUU | Abneutral | 0.1223 | 32 | 31 | 0.7289703 |
| CHG | TE |  |  |  | ABselectUU | Abneutral | 0.0759 | 32 | 31 | 0.7847896 |
| CHG | global |  |  |  | ABselectMM | Abneutral | 0.0312 | 32 | 31 | 0.8609342 |
| CHG | exon |  |  |  | ABselectMM | Abneutral | 0.0000 | 32 | 31 | 1.00E+00 |
| CHG | promoter |  |  |  | ABselectMM | Abneutral | 0.1063 | 32 | 31 | 0.7466025 |
| CHG | TE |  |  |  | ABselectMM | Abneutral | 0.0779 | 32 | 31 | 0.7820306 |
| CHH | global | NA | NA | NA | ABneutral | Abnull | 0.6695 | 35 | 31 | 0.6180636 |
| CHH | exon | NA | NA | NA | ABneutral | Abnull | 0.5049 | 35 | 31 | 0.732369 |
| CHH | promoter | NA | NA | NA | ABneutral | Abnull | 1.2939 | 35 | 31 | 0.2938961 |
| CHH | TE | NA | NA | NA | ABneutral | Abnull | 0.3691 | 35 | 31 | 0.8287454 |
| CHH | global |  |  |  | ABselectUU | Abneutral | 0.0018 | 32 | 31 | 0.9661093 |
| CHH | exon |  |  |  | ABselectUU | Abneutral | 0.0000 | 32 | 31 | 1 |
| CHH | promoter |  |  |  | ABselectUU | Abneutral | 0.0016 | 32 | 31 | 0.9687817 |
| CHH | TE |  |  |  | ABselectUU | Abneutral | 0.7337 | 32 | 31 | 0.3982717 |
| CHH | global |  |  |  | ABselectMM | Abneutral | 0.0079 | 32 | 31 | 0.9296158 |
| CHH | exon |  |  |  | ABselectMM | Abneutral | 0.0000 | 32 | 31 | 1 |
| CHH | promoter |  |  |  | ABselectMM | Abneutral | 0.0003 | 32 | 31 | 0.9871071 |
| CHH | TE |  |  |  | ABselectMM | Abneutral | 0.0325 | 32 | 31 | 0.8581938 |

FM = Full model RM = Reduced model df = degrees of freedom Best performing model

**Table S4:** Epimutation rate estimates and model selection results for pedigree MA3.

| context | annotation | alpha | beta | beta/alpha | FM | RM | F-value | df RM | df FM | P--value |
| --- | --- | --- | --- | --- | --- | --- | --- | --- | --- | --- |
| CG | global | 0.0001942304 | 0.0008340002 | 4.294 | ABneutral | Abnull | 136.2307 | 65 | 61 | 1.103063E-29 |
| CG | exon | 0.0005635082 | 0.001829657 | 3.247 | ABneutral | Abnull | 210.1967 | 65 | 61 | 6.177088E-35 |
| CG | promoter | 0.0001128601 | 0.001141507 | 10.114 | ABneutral | Abnull | 108.4535 | 65 | 61 | 5.190251E-27 |
| CG | TE | NA | NA | NA | ABneutral | Abnull | 1.0608 | 65 | 61 | 0.383711 |
| CG | global |  |  |  | ABselectUU | Abneutral | 0.0000 | 62 | 61 | 1 |
| CG | exon |  |  |  | ABselectUU | Abneutral | 0.0000 | 62 | 61 | 1 |
| CG | promoter |  |  |  | ABselectUU | Abneutral | 0.1053 | 62 | 61 | 0.7467267 |
| CG | TE |  |  |  | ABselectUU | Abneutral | 0.0000 | 62 | 61 | 1 |
| CG | global |  |  |  | ABselectMM | Abneutral | 0.0001 | 62 | 61 | 0.9934013 |
| CG | exon |  |  |  | ABselectMM | Abneutral | 0.0000 | 62 | 61 | 1 |
| CG | promoter |  |  |  | ABselectMM | Abneutral | 0.0000 | 62 | 61 | 1 |
| CG | TE |  |  |  | ABselectMM | Abneutral | 0.1116 | 62 | 61 | 0.739439 |
| CHG | global | NA | NA | NA | ABneutral | Abnull | 2.0182 | 65 | 61 | 0.1030624 |
| CHG | exon | NA | NA | NA | ABneutral | Abnull | 0.0141 | 65 | 61 | 0.9995992 |
| CHG | promoter | 1.783862E-06 | 0.0001055335 | 59.160 | ABneutral | Abnull | 5.0076 | 65 | 61 | 0.001479819 |
| CHG | TE | 5.517707E-05 | 0.0001307905 | 2.370 | ABneutral | Abnull | 5.7078 | 65 | 61 | 0.0005720866 |
| CHG | global |  |  |  | ABselectUU | Abneutral | 0.9363 | 62 | 61 | 0.3370451 |
| CHG | exon |  |  |  | ABselectUU | Abneutral | 0.0000 | 62 | 61 | 1 |
| CHG | promoter |  |  |  | ABselectUU | Abneutral | 0.0707 | 62 | 61 | 0.7912417 |
| CHG | TE |  |  |  | ABselectUU | Abneutral | 0.5221 | 62 | 61 | 0.4727021 |
| CHG | global |  |  |  | ABselectMM | Abneutral | 0.8975 | 62 | 61 | 0.3471843 |
| CHG | exon |  |  |  | ABselectMM | Abneutral | 0.0000 | 62 | 61 | 1.00E+00 |
| CHG | promoter |  |  |  | ABselectMM | Abneutral | 0.0000 | 62 | 61 | 1 |
| CHG | TE |  |  |  | ABselectMM | Abneutral | 0.8804 | 62 | 61 | 0.3517958 |
| CHH | global | NA | NA | NA | ABneutral | Abnull | 1.0187 | 65 | 61 | 0.404864 |
| CHH | exon | NA | NA | NA | ABneutral | Abnull | 0.1767 | 65 | 61 | 0.9495845 |
| CHH | promoter | 1.71043E-08 | 4.1274E-06 | 241.308 | ABneutral | Abnull | 2.9878 | 65 | 61 | 0.02558529 |
| CHH | TE | NA | NA | NA | ABneutral | Abnull | 0.2803 | 65 | 61 | 0.8896153 |
| CHH | global |  |  |  | ABselectUU | Abneutral | 0.7073 | 62 | 61 | 0.4036315 |
| CHH | exon |  |  |  | ABselectUU | Abneutral | 0.4033 | 62 | 61 | 0.5277759 |
| CHH | promoter |  |  |  | ABselectUU | Abneutral | 0.0000 | 62 | 61 | 1 |
| CHH | TE |  |  |  | ABselectUU | Abneutral | 0.0000 | 62 | 61 | 1 |
| CHH | global |  |  |  | ABselectMM | Abneutral | 1.2129 | 62 | 61 | 0.275082 |
| CHH | exon |  |  |  | ABselectMM | Abneutral | 0.2588 | 62 | 61 | 0.6127902 |
| CHH | promoter |  |  |  | ABselectMM | Abneutral | 0.0000 | 62 | 61 | 1 |
| CHH | TE |  |  |  | ABselectMM | Abneutral | 0.0684 | 62 | 61 | 0.7945567 |

FM = Full model RM = Reduced model df = degrees of freedom Best performing model

**Table S5:**
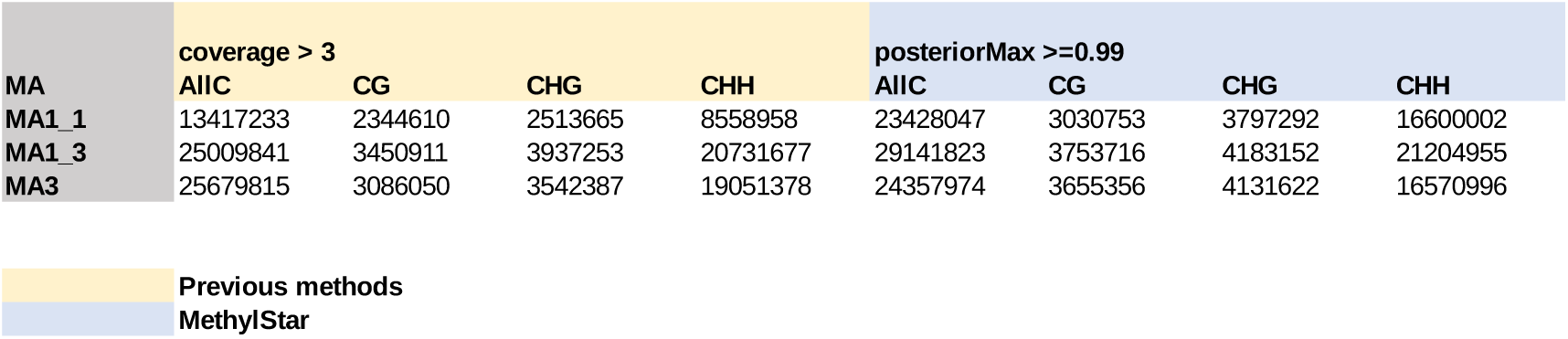
Pre-processing of WGBS data using MethylStar increases the number of high-confident cytosines that can be used for epimutation analysis compared with previous pre-processing approaches.

